# Development of circulating cell-free DNA reduced representation bisulfite sequencing for clinical methylomics diagnostics

**DOI:** 10.1101/663195

**Authors:** Andries De Koker, Lennart Raman, Ruben Van Paemel, Malaïka Van der Linden, Sofie Van de Velde, Paul van der Leest, Bram Van Gaever, Anxhela Zaka, Karim Vermaelen, Ingel Demedts, Veerle Surmont, Ulrike Himpe, Franceska Dedeurwaerdere, Liesbeth Ferdinande, Kathleen Claes, Jilke De Wilde, Mieke Zwager, Vincent D. de Jager, T. Jeroen N. Hiltermann, Bram De Wilde, Björn Menten, Katleen De Preter, Ed Schuuring, Jo Van Dorpe, Nico Callewaert

**Author notes:** N.C. and J.V.D. share corresponding authorship. and. A.D.K. and L.R. share first authorship.

## Abstract

Methylation profiling of circulating cell-free DNA (cfDNA) is of great interest as a liquid biopsy assay for the detection and monitoring of cancer and other pathologies. Here we describe ‘circulating cell-free DNA reduced representation bisulfite sequencing’ (cf-RRBS), enabling the use of highly effective RRBS on fragmented plasma cfDNA. This method enriches the CpG-rich RRBS target regions by enzymatic degradation of all off-target DNA rather than by targeted capture, in contrast to previous methods. Critical steps are fully enzymatic in a single-tube, making it rapid, cost-effective, robust, and easily implemented on a liquid-handling station for high-throughput sample preparation. We benchmark cf-RRBS results to those obtained by previous more complex methods and exemplify its use for accurate non-invasive subtyping of lung cancer, a frequent onco-pathology task. cf-RRBS enables any molecular pathology lab to tap into the cfDNA methylome, only making use of off-the-shelf reagents and open-source data analysis tools.

**One Sentence Summary:** Novel methodology enables facile and cost-effective methylation profiling of fragmented plasma DNA, allowing for routine liquid biopsy clinical diagnostics, exemplified here for differential diagnosis of lung cancer subtypes.

## INTRODUCTION

Altered DNA methylation is a near universal hallmark of oncogenic transformation and, in a broader sense, of tissue pathology (1). Recent studies (2–5) have demonstrated that methylation profiles derived from circulating cell-free DNA (cfDNA) not only enable tumor detection, but also reveal the tissue of origin for various cancer entities. Detection of altered DNA methylation patterns through DNA methylation based diagnostics in easily sampled body fluids like blood plasma, i.e. through ‘liquid biopsy’, may be used to detect and classify cancer.

The most popular tissue methylation profiling approaches are based on the array technology of Illumina (Infinium 450K or EPIC BeadChip) (6,7) or on Reduced Representation Bisulfite Sequencing (RRBS) (8), which fail on highly fragmented low input DNA such as cfDNA in blood. At present, whole-genome bisulfite sequencing (WGBS) on cfDNA can be used, but remains too costly for routine application, mainly because of its non-targeted nature, requiring a substantial sequencing budget in order to enable robust quantitation of methylation levels (9). Hence, the key to cost-effectively obtaining robust information on the methylation status of regulatory sections of the genome leaked by dying cells as circulating cell-free plasma DNA, is not sequencing the entire genome, but to focus on CpG-rich genomic regions (10). Therefore, methods to capture CpG-rich regions have been developed, using expensive custom-designed and often proprietary reagents, such as capture probes in SeqCap Epi (11) and similar products, or antibodies binding to methylated DNA in cfMeDIP-seq (12). These methods all involve highly complex and lengthy protocols that are challenging to automate (Figure 1A).

**Figure 1.**
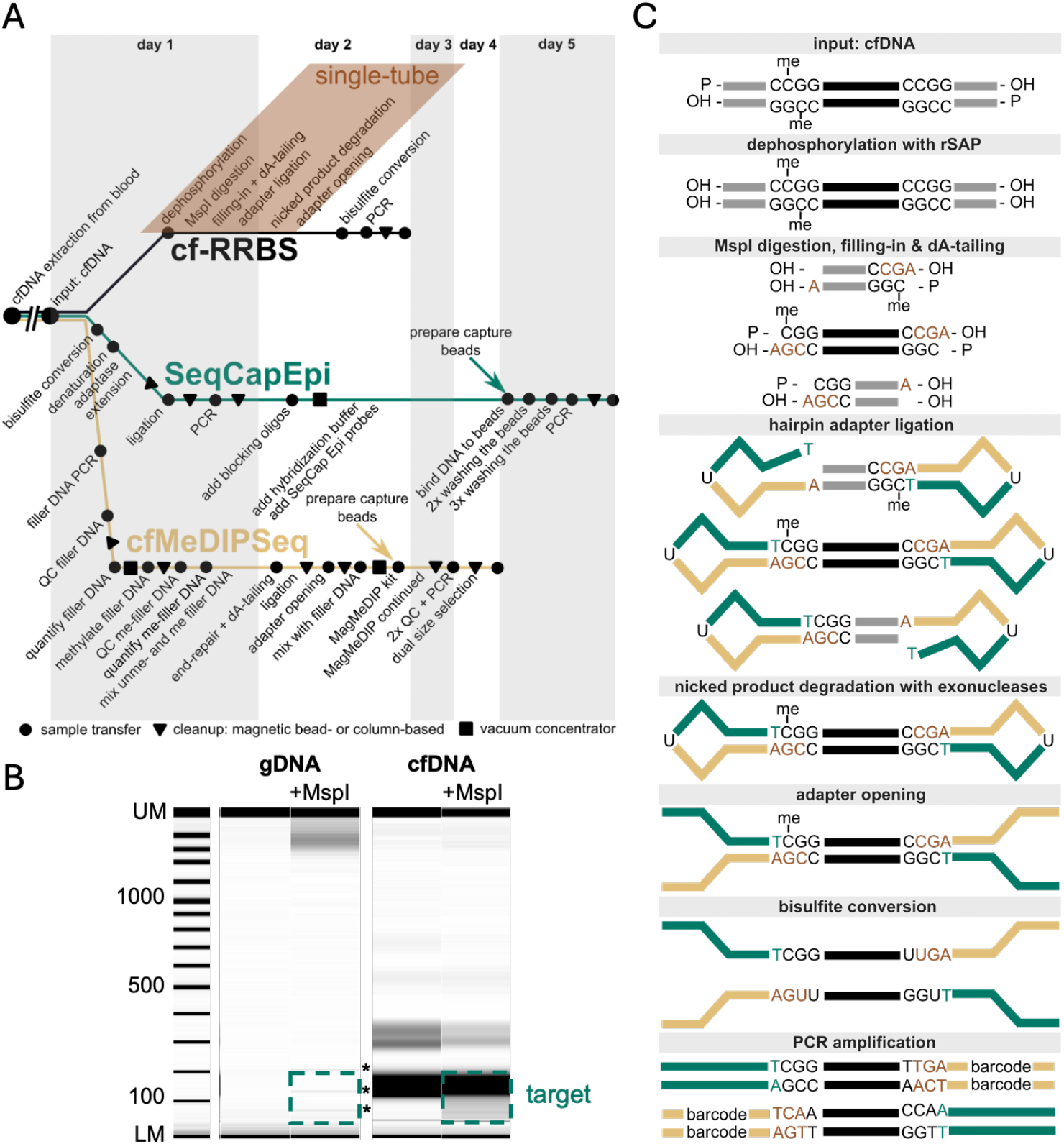
cf-RRBS enables reduced representation bisulfite sequencing on highly fragmented DNA. **(A)** A simplified protocol comparison between cf-RRBS, SeqCap Epi and cfMeDIPSeq in function of time. cf-RRBS is technically much less complex than target capture-based and antibody capture-based methods. Both SeqCap Epi and cfMeDIPSeq require a much more complex sequence of sample processing steps than cf-RRBS, which also takes much less time. Additional steps include sample transfers (dots), solid-phase reversible immobilization magnetic bead purifications (triangles), vacuum concentration steps (rectangles), many of which are not easily amenable to automation. (B) A digital gel image of gDNA and cfDNA, either undigested or MspI-digested. In classic RRBS, MspI/MspI-fragments ranging from 20 to 200 bp (indicated with a green rectangle; ‘the target’) are to be extracted from the gel. The undigested gDNA runs above the upper marker (UM) and is not observed. After MspI digestion, three (indicated with asterisks) characteristic satellite DNA bands appear near the lower maker (LM) and this region contains the target MspI/MspI short fragments that are then gel-extracted to achieve the enrichment that results in the reduced representation of the genome. With cfDNA, such reduced representation target extraction is impossible, since cell-free fragments are naturally highly fragmented and the size range completely overlaps with the target MspI/MspI fragment size range. (C) A simplified cf-RRBS workflow, where thick lines indicate DNA fragment strands. Fragments are subject to dephosphorylation, MspI digestion, dA-tailing, and ligation of hairpin-shaped adapters. Only MspI/MspI-fragments yield unnicked DNA molecules without free ends, providing protection against exonucleases, which degrade all off-target DNA molecules, hence achieving reduced representation target enrichment even if the starting DNA is highly fragmented, such as in plasma cfDNA and in FFPE-tissue extracted DNA.

Classical RRBS (8), in contrast, achieves its CpG-rich region focus by digesting high molecular weight (high-MW) genomic DNA (gDNA) extracted from tissues with a restriction enzyme (typically MspI) that cuts CpG-rich sequences (i.e., ‘CpG islands’). This is followed by size-based gel electrophoresis separation of the short MspI/MspI-fragments resulting from the genomic DNA input material, empowering high sequencing coverage of the CpG-enriched ∼2% of the genome within these short MspI/MspI fragments. This ∼2% of the genome has been amply demonstrated as a very effective measure of genome-wide methylation status ever since RRBS was invented (8,13). The required sequencing costs are similar to what is presently routinely used in Non-Invasive Prenatal Testing (NIPT) (14), which serves as a benchmark for clinical implementation, as it is one of the most widespread cfDNA sequencing-based molecular diagnostic tests in use today. RRBS does not require any customized target-selective reagents, making it a universal method for genome-wide methylome profiling across pathologies. While these properties conceptually make RRBS attractive for its use in liquid biopsies, its current implementations are not suitable for this purpose, as the minute amounts of cfDNA input are naturally (highly) fragmented, precluding the required size separation of the MspI/MspI fragments from the fragmented input cfDNA (Figure 1B).

To overcome this, we set out to develop cell-free RRBS (cf-RRBS), which incorporates key RRBS innovations to fit the fragmented cfDNA input material. At the same time, we removed low-throughput manual steps such as gel separation, enabling high-throughput automation that is essential for robust use in diagnostic molecular pathology laboratories.

Here we report on the development of cf-RRBS, characterization and benchmarking of the methodology, and we illustrate its power in a key application: non-invasive histology subtyping of lung cancer.

## RESULTS

### cf-RRBS method development

Conceptually, rather than affinity-purifying a CpG-dense genomic target subsample as in previous methods, we inverted the problem, and chose to enzymatically destroy all off-target DNA in the sample upon protecting only the targeted MspI/MspI restriction-site flanked CpG-rich genomic subsample.

In short (Figure 1C), in cf-RRBS, input DNA is first dephosphorylated by recombinant Shrimp Alkaline Phosphatase (rSAP). Fragments are subsequently digested by MspI, resulting in novel phosphorylated 5’-ends, marking MspI cuts. Upon 5’ overhang filling, dA-tailing and hairpin-shaped adapter ligation, exclusively MspI/MspI-fragments are typified by fully ligated and hence unnicked ends. These ‘circular’ (i.e., with hairpins at both ends) MspI/MspI-fragments are uniquely resistant to an optimized mixture of single- and double-strand-specific 3’>5’ and/or 5’>3’ exonucleases, whereas all other DNA is degraded, effectively achieving the RRBS target enrichment. Finally, following adapter opening and bisulfite conversion, fragments are amplified, and a sequencing library is obtained. With robustness and ease of automation in mind, we thoroughly optimized the implementation of this method concept, such that only sequential reagent additions and incubations are needed in a single tube, up to the first DNA amplification. This required the reagent concentrations and compositions to be precisely tuned to avoid interference with subsequent enzymatic reactions: e.g., dATP is used by Klenow for dA-tailing, but an excessive amount inhibits T4 DNA ligase during hairpin ligation (15). Similarly, heat-labile variants of selected enzymes were introduced, including rSAP and Antarctic thermolabile Uracil-DNA Glycosylase (used for adapter opening), such that enzyme inactivation can simply be achieved with an intermittent heating step.

The elimination of all intermittent purification steps and tube transfers up to bisulfite conversion allowed for robustness in working with the few nanograms of DNA extractable from patient plasma which is critically important for robust clinical utility (Figure 1A). Other design optimization parameters included short runtime and low cost, achieving a total cost, including sequencing, not surpassing that of the routinely used NIPT test, a benchmark for successful cfDNA sequencing-based molecular diagnostic tests. All of this required substantial optimization and we recommend users to adhere tightly to the finalized protocol parameters set out in detail in the Online Methods, which also includes how to automate cf-RRBS on a TECAN EVO200 liquid handling platform.

### cf-RRBS target enrichment performance

Most cfDNA fragments have a length of one nucleosomal winding (i.e. 166bp), with a smaller fraction corresponding to the length of DNA wound around two nucleosomes, and vanishingly small amounts of multiple-nucleosome fragments (16). Hence, we defined the target genomic subsample of cf-RRBS as all MspI/MspI-fragments sized 20-200 bp, as longer MspI/MspI genomic fragments would only rarely be present. These targeted MspI/MspI fragments make up ∼2% of the human genome, consistent with the 42.4 +/− 3.2-fold enrichment of this target observed with the cf-RRBS method (n=16) (Figure 2A), resulting in >90% of the reads mapping to the target MspI/MspI regions (Figure 2B). In Figure 2A we also illustrate that the >40-fold target enrichment factor achieved on the highly fragmented cfDNA is in the same range as is achieved with gel-separation based RRBS on high-MW gDNA. The data also indicate that this gel-based size-separation classical RRBS implementation is indeed incapable of achieving high and robust target enrichment when applied to fragmented cfDNA.

**Figure 2.**
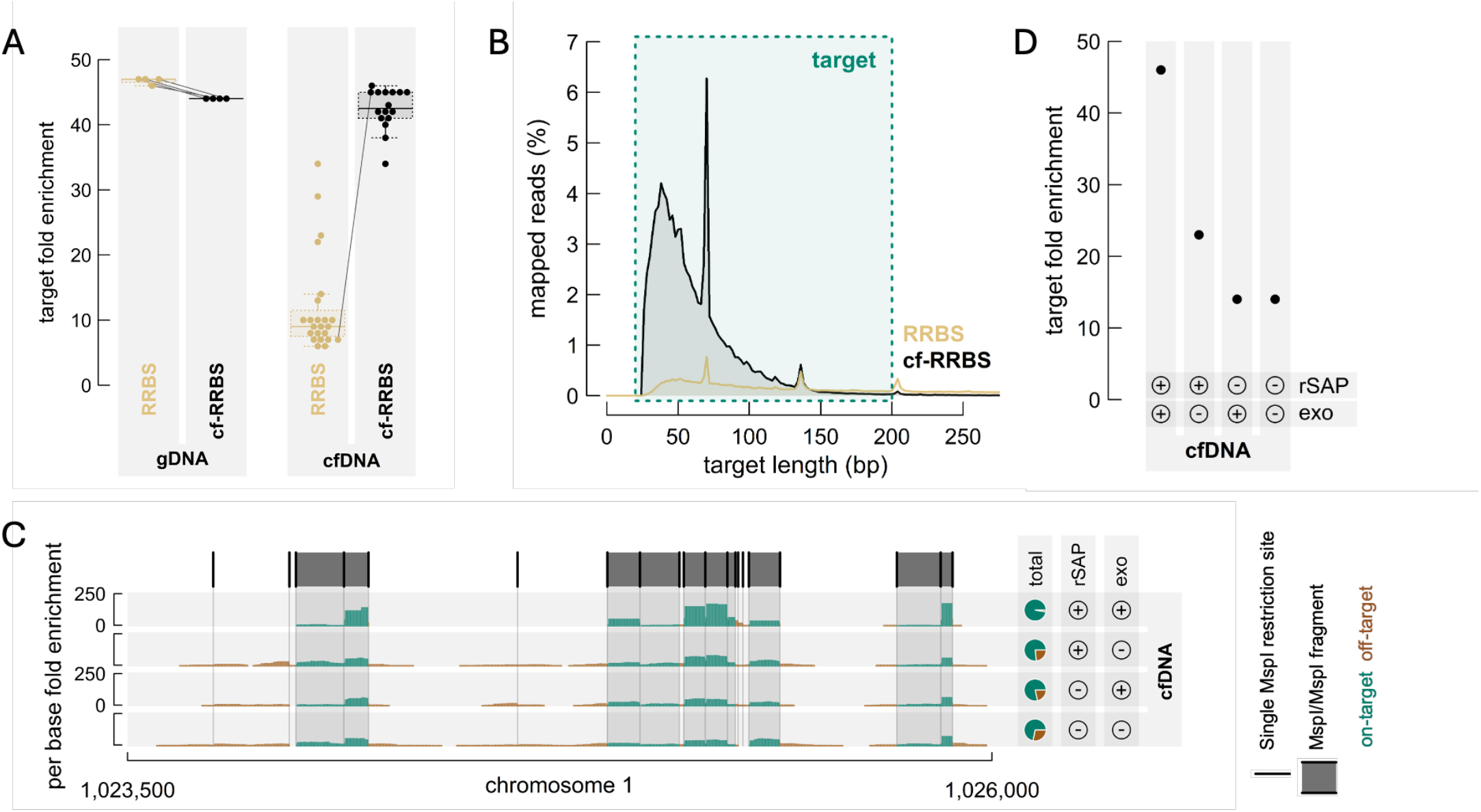
cf-RRBS solves the target enrichment problem of classic RRBS on cfDNA. **(A)** While a similar target fold enrichment is observed between classic RRBS and cf-RRBS on gDNA (n=4), RRBS demonstrates poor enrichment when executed on cfDNA (n=23), which however can be achieved effectively by using the cf-RRBS method (n=16). Same-sample dots (paired samples) are connected. (B) A histogram of the mapped reads in function of the target length shows that cf-RRBS reads map the target at a much higher frequency than classic RRBS. (C) The per base fold enrichment of a representative chromosomal stretch of 2.5 kb is visualized across four samples, annotated by cf-RRBS protocol control deviations at the right-hand side, including intentionally missing steps concerning rSAP dephosphorylation, exonuclease (exo) degradation, both or none illustrating that all steps in the cf-RRBS method contribute to its efficiency at enriching the MspI/MspI fragments and depleting the DNA that is not contained in such fragments. The top track presents *in silico* predicted MspI digestion sites and MspI/MspI-fragments (sized 20-200 bp). Pie charts clarify on- and off-target fractions of the visualized genomic stretch. (D) Similar to C but here the overall genome-wide target fold enrichment is shown instead of for the randomly selected genomic stretch of panel C.

Two key steps in the cf-RRBS method enable specific MspI/MspI-fragment enrichment without relying on DNA size: first, marking of natural fragment ends through dephosphorylation by rSAP, enabling bistranded ligation of closed hairpin adapters only to MspI/MspI digested target fragments (which regain 5’ phosphates at both ends in the digest); and, second, degrading unprotected off-target fragments by a mixture of exonucleases (Figure 1C). We evaluated the effect of omitting one or both of these steps. As illustrated by visualizing the resulting sequencing reads that map to a randomly chosen 2.5 kb genomic stretch on chromosome 1, removing either the initial dephosphorylation or the exonuclease digestion of off-target DNA indeed obviates the precise enrichment of MspI/MspI-fragments (Figure 2C). These observations were confirmed at genome-wide scale (Figure 2D): only with inclusion of both key steps can the >40-fold MspI/MspI target enrichment be achieved.

### cf-RRBS quantifies methylation status highly reproducibly and yields equivalent results with established more complex and/or more costly cfDNA methylation profiling methods

The methylation profiling accuracy of cf-RRBS was benchmarked by pairwise concordance analyses using cfDNA samples that were also analyzed by SeqCap Epi (SCE, genome-wide targeted capture method from Roche-Nimblegen). Both CpG site- and island-level ‘beta values’ (i.e., the ratio of the number of reads that indicate methylation over the total number of reads mapping to a CpG site or CpG island) were considered, measuring agreement by Pearson correlation and mean absolute error (∼the average ‘disagreement’ between both methods). The genomic regions targeted by SeqCap Epi and cf-RRBS are only partially overlapping, so these analyses are limited to the genomic regions that are sampled by both technologies. Results from a representative plasma sample are shown (Figure 3A).

**Figure 3.**
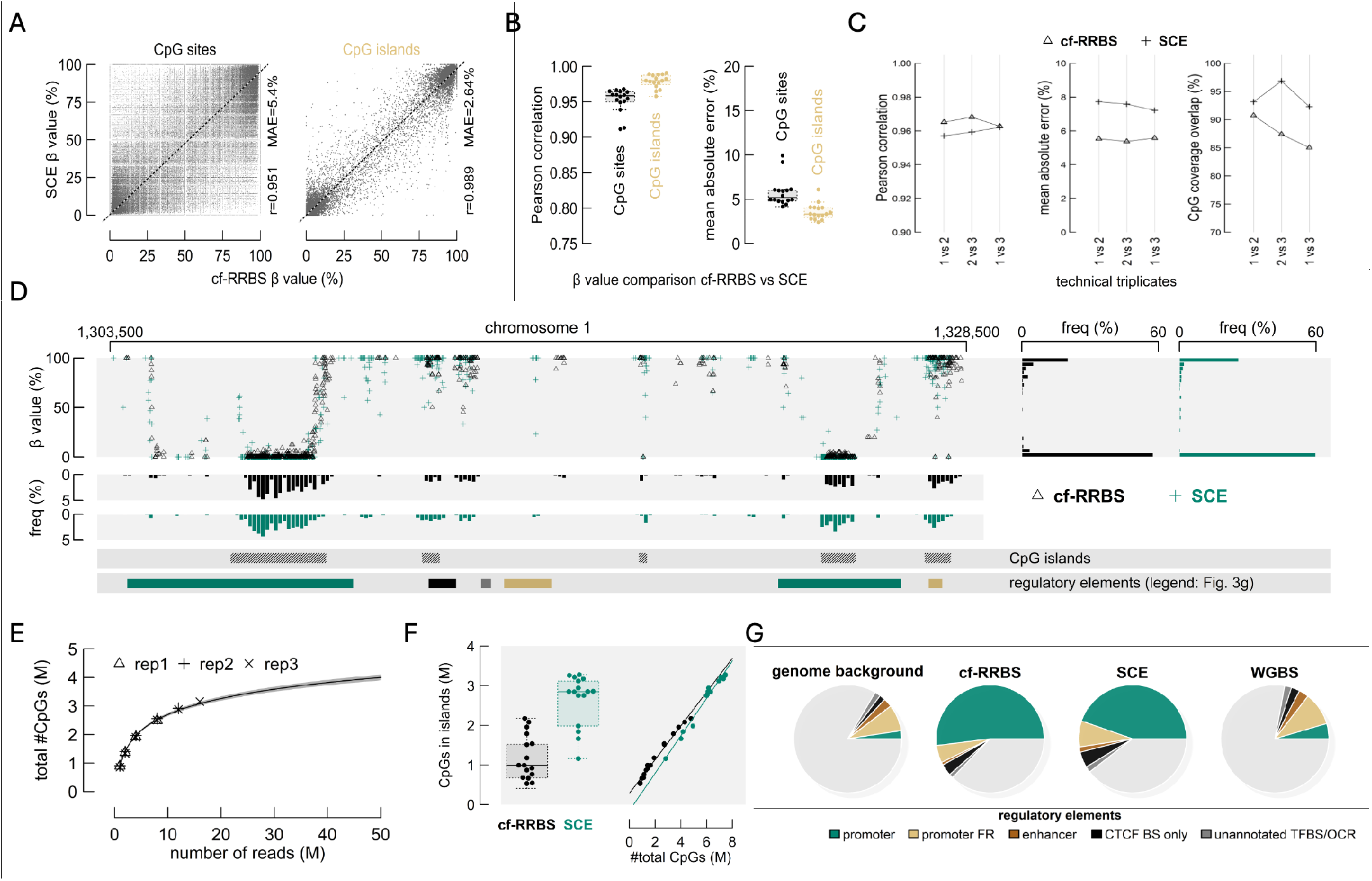
Methylation profiling with cf-RRBS is accurate. **(A)** Methylation profile scatter plot comparison between cf-RRBS and SeqCap Epi of representative patient LP10’s cfDNA, at CpG site-level (left) and at CpG island-level (right). Least squares fit (dotted line), corresponding Pearson correlation (r), and the mean absolute error (MAE) are given for both comparisons. (B) Swarm/boxplot combinations of the Pearson correlation (left) and mean absolute error (right) between cfDNA samples analyzed by cf-RRBS and SeqCap Epi. (C) Pairwise comparisons between cfDNA samples analyzed in triplicate by cf-RRBS and SeqCap Epi. Paired relations are connected. Analyses include the Pearson correlation between CpG site beta values (left); the mean absolute error between CpG site beta values (middle); and the positional overlap in CpG calls (right), calculated as the number of recurrent CpGs over the total number of CpGs in the sample with the lowest sequencing depth. (D) Methylation profiles of representative patient LP10’s cfDNA derived by cf-RRBS (black) and SeqCap Epi (green) of a genomic stretch sized 25 kb. Frequency histograms of CpG beta values are shown, one for each technique (right). Similarly, the spatial distribution of CpG calls is clarified by positional frequency histograms below. The bottom two bars contain tracks of CpG islands and regulatory elements, see legend of panel G. (E) the number of raw cfDNA sequencing reads is shown in relation to the total number of CpG calls for three technical cf-RRBS replicates, randomly subsampled to 1, 2, 4, 8, 12 and 16 million reads. A log regression fit, with 95% confidence interval (given by shading), indicates three million CpG calls are obtained with less than 20 million reads. (F) Swarm/boxplot combinations of the number of CpG calls located in CpG islands compared between cfDNA samples analyzed by cf-RRBS and SeqCap Epi. The scatter plot at the right-hand side provides the relation with the total number of CpG calls for each technique, given by least squares fits. cf-RRBS provides relatively more calls located in CpG islands. (G) Pie charts indicate the average fractions of CpGs that cover regulatory elements per cfDNA methylation profiling method. The far-left pie demonstrates the genomic background. Included regulatory elements are unannotated transcription factor binding sites (TFBSs) and open chromatin regions (OCRs); (parts of) CTCF binding sites (BSs) that do not travel across other elements; enhancers; promoter flanking regions (FRs); and promoters.

Correlations were high between both methods: average r = 0.96 for sites and r = 0.98 for islands, and mean absolute errors were consistently below 6% (Figure 3B). Of note, both target-enrichment methods compared equally well to results from WGBS analysis: concordance with WGBS was substantially better at the CpG island level (which averages the methylation status of many CpG sites) than at individual CpG level, which is likely caused by inaccurate single CpG methylation status sequencing by the low per-site coverage that is obtained in WGBS (Figure S1).

A selected cfDNA sample was analyzed in triplicate by both cf-RRBS and SeqCap Epi to estimate technical variation. Both methods performed highly consistently (r>0.95; Figure 3C). The CpG site call ‘overlap’ between technical replicates was more than 80% and was largely similar for both methods (Figure 3C, right). In exploratory experiments, and in keeping with the method having been designed to operate on highly fragmented DNA, we have also successfully applied cf-RRBS to DNA extracted from formalin-fixed paraffin-embedded (FFPE) tissues and report on this elsewhere (17). Hence, the same cf-RRBS method can be used both on patient cfDNA and on the most commonly available archived samples of diseased tissues in histopathological collections, which should greatly facilitate making direct correlations between cfDNA methylation alterations and those in the diseased tissues under study (which can be used as reference methylome atlases to deconvolute cf-RRBS data obtained from plasma).

### cf-RRBS predominantly targets gene promoters

The functional genomic coverage of cf-RRBS was compared to method alternatives by mapping individual CpG calls to CpG islands and publicly available tracks of regulatory elements (i.e., promoters; promoter flanking regions; enhancers; CTCF (CCCTC-Binding factor) binding sites; and remaining unannotated ChIP-Seq transcription factor binding sites and DNase I hypersensitivity open chromatin regions). A representative 25 kb chunk of the methylation profiles of the same cfDNA sample, obtained either by cf-RRBS or by SeqCap Epi, visually demonstrates that the spatial distribution of CpG calls is, as expected, highly enriched in CpG islands and regulatory elements (Figure 3D).

The number of CpG methylation calls is of course highly related to sequencing depth. At 20 million reads (2 x 75 bp), cf-RRBS provides methylation information on more than 3 million CpG sites (Figure 3E). Since the SeqCap Epi kit was designed by the manufacturer to capture-enrich a larger genomic subsample than cf-RRBS, it naturally covers more CpG sites when this target is sequenced at a depth according to protocol guidelines, requiring about 50 million reads (2 x 100 bp). However, the fraction (= CpG calls in islands/total CpG calls) of CpGs calls in islands is, overall, greater for cf-RRBS than for SeqCap Epi (Figure 3F). In contrast to WGBS, both of the CpG-rich genomic area-targeting methods have a much higher CpG call rate in annotated regulatory elements of the genome (Figure 3G). Notably, 53.4% of CpG calls made by cf-RRBS cover promoters, whilst these, as currently annotated in the human genome, only embody ∼2.5% of the human genome (18). Differential promoter methylation is a key information source as to the cell of origin of methylated DNA and hence of key diagnostic importance (19).

### Non-invasive cfDNA methylation profiling by cf-RRBS allows specific histological lung cancer subtyping

While cf-RRBS is widely applicable and studies are ongoing in our lab and several others in a multitude of clinical diagnostic settings (2, 17, 20, 21), we illustrate its utility in the present original method invention paper by providing a proof-of-principle study on lung cancer. Lung cancer is fundamentally subdivided in small cell lung cancer (SCLC) and non-small cell lung cancer (NSCLC). Approximately 85% of diagnoses are NSCLC, of which the most common subtypes are adenocarcinoma (LUAD) and squamous cell carcinoma (LUSC). These distinctions have long been therapeutically relevant, especially for targeted treatment approaches (22) and even for chemotherapy selection (23,24). Most importantly, diagnosing SCLC currently suffices to initiate (chemo)therapy, as for now no oncogene-targeted treatments are routinely used, rendering further molecular characterization as yet unnecessary. Also, transformation to SCLC is seen as a mechanism of treatment resistance and relapse upon treatment of EGFR-mutated NSCLC (25), an event which might be intercepted using liquid biopsy-based disease monitoring instead of invasive tissue biopsy.

We analyzed a first cohort, obtained from Ghent University Hospital (UZ Gent) and from the AZ Delta Hospital Roeselare, of 51 liquid biopsies (lung cancer n=42 and non-tumor pulmonary disease n=9) by cf-RRBS. A second cohort, obtained from the University Medical Center of Groningen (UMCG), of 63 liquid biopsies (lung cancer n=33 and healthy controls n=30) was analyzed as well by cf-RRBS, for external classifier validation purposes. Clinical parameters per cohort are shown in Table 1.

**Table 1:** Clinical parameters per cohort.

| Patient cohort UZ Gent |  |  |
| --- | --- | --- |
| Variable | Parameter | No. of patients |
| Age | Median (range) | 66 (44-80) |
| Sex | Male | 30 (71%) |
|  | Female | 12 (29%) |
| Smoking history | Smoker | 14 (33%) |
|  | Ex-smoker | 25 (60%) |
|  | Non-smoker | 2 (5%) |
|  | unknown | 1 (2%) |
| Stage | III | 6 (14%) |
|  | IV | 36 (86%) |
| Histology | Adenocarcinoma | 18 (43%) |
|  | Squamous cell carcinoma | 12 (29%) |
|  | Small cell | 12 (29%) |

| Control cohort UZ Gent |  |  |
| --- | --- | --- |
| Variable | Parameter | No. of patients |
| Age | Median (range) | 59 (41-71) |
| Sex | Male | 7 (78%) |
|  | Female | 2 (22%) |

| Patient cohort UMCG |  |  |
| --- | --- | --- |
| Variable | Parameter | No. of patients |
| Age | Median (range) | 64 (40-83) |
| Sex | Male | 15 (45%) |
|  | Female | 18 (55%) |
| Smoking history | Smoker | 24 (73%) |
|  | Ex-smoker | 8 (24%) |
|  | Non-smoker | 1 (3%) |
|  | unknown | 0 (0%) |
| Stage | III | 2 (6%) |
|  | IV | 31 (94%) |
| Histology | Adenocarcinoma | 22 (67%) |
|  | Squamous cell carcinoma | 11 (33%) |
|  | Small cell | 0 (0%) |

| Control cohort UMCG |  |  |
| --- | --- | --- |
| Variable | Parameter | No. of patients |
| Age | Median (range) | 63 (48-77) |
| Sex | Male | 14 (47%) |
|  | Female | 16 (53%) |

A customized data processing and modeling approach for the methylation-based tumor classification was developed. First, we clustered the CpGs of which the methylation level could be determined in >50% of the UZGent healthy control cfDNA samples (n = 46; 19 males and 27 females, median age 31 (23-61)), according to a clustering scheme (detailed in the Supplementary Methods) that optimally accommodates that cf-RRBS-detected CpGs are located close together in short MspI/MspI fragments. This resulted in more than 500,000 such clusters (Table S3A), and methylation levels on these clustered CpGs were averaged into a cluster-specific beta value (i.e. methylation fraction). Unstable clusters were removed (i.e. standard deviation beta value >1%) within this set of healthy control cf-RRBS datasets (n = 46), in this way minimizing interference of e.g. age-or sex-related methylation alterations (26,27). This resulted in a set of 28,128 of such CpG clusters with stable methylation in healthy controls. Illustrating how publicly available tumor methylation data can be reutilized in designing classification models for cf-RRBS-based diagnosis, we extracted these same CpG-cluster methylation values from 682 lung tumor methylomics datasets, enabling us to identify those most informative for the lung tumor subtype classification task at hand. This procedure resulted in 473 clusters that were propagated in our classification models (Table S3B). t-distributed Stochastic Neighbor Embedding (t-SNE) analysis using the selected clusters demonstrated an almost complete separation of the different lung cancer subtypes in the publicly sourced tumor methylation reference dataset (Figure 4A). Hence, the prioritized set of CpG clusters was used to perform deconvolution (5), designed to assign cf-RRBS data of patients to four subfractions: healthy, LUAD, LUSC and SCLC.

**Figure 4.**
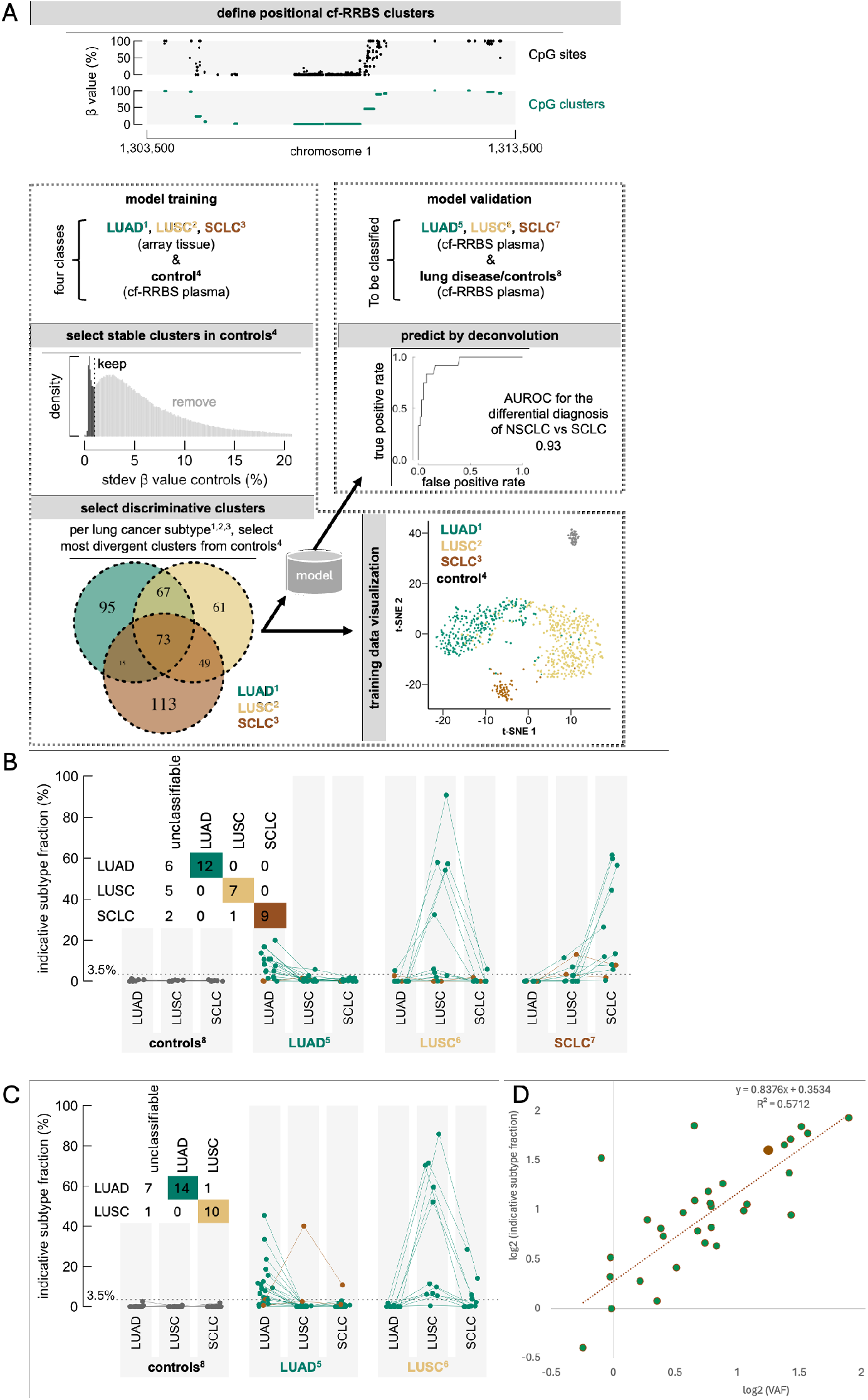
Methylation profiles obtained by cf-RRBS enable non-invasive lung cancer subtyping at high specificity. **(A)** Predictive modeling workflow. Prior to modeling, CpG clusters according to spatial distribution optimized for cf-RRBS were defined in order to reduce data complexity with minimal data loss. Control liquid biopsies (n=46); and public methylation datasets of LUAD, LUSC and SCLC tissue samples (n=682), served as a reference set. Based on these, CpG clusters for modeling were selected using a two-step procedure: first, variable clusters in controls were removed based on the standard deviation (stdev); second, per lung cancer subtype, the most divergent clusters in comparison to healthy donor plasma cfDNA controls were kept (cluster overlap between subtypes is indicated by Venn diagram). The reference set is of lung cancer tissue methylation datasets and healthy donor’s cfDNA cf-RRBS visualized by t-SNE analysis, based on these selected clusters. Finally, a linear deconvolution model for the cfDNA methylation ß-values in these clusters was trained, attributing fractions of the cfDNA to the four deconvoluted entities such that the linear combination of each reference ß-values multiplied by the attributed fractions results in the observed cfDNA ß-value. Following modeling by deconvolution, validation was executed using two cohorts, patient (n=42 or n=33) and control liquid biopsies (n=9 or n=30). Differential diagnosis of NSCLC versus SCLC across both cohorts is indicated by receiver operating characteristic analysis, summarized by AUROC. (B-C) Indicative subtype fractions and confusion table at the 3.5% ‘noise’ cutoff of each lung cancer subtype for in-house cohort and (B) and UMCG cohort (C). Samples are grouped per type. For controls and patients, every fraction is shown, where same-sample dots are connected. The dotted line at 3.5% represent a ‘noise’ cutoff: control liquid biopsies are consistently below this limit. (D) Correlation log2(VAF) versus log2(indicative subtype fraction) on UMCG cohort. Correct prediction (green) and wrong prediction (brown).

This model was then used to classify the cf-RRBS-profiled patients and controls according to the model subgroup for which the highest disease-specific fraction was assigned. The multi-class area under the curve (AUC) across both cohorts to differentiate the three lung cancer classes and controls was 0.96. A receiver operating characteristic (ROC) curve is shown differentiating control versus tumor in Fig S2. For differentiation of lung cancer subtypes in patients with a predetermined presence of a tumor, the multi-class AUC is 0.96. For treatment purposes, the differentiation of NSCLC and SCLC is most important; the ROC curve using the SCLC score with an AUC of 0.93 is shown in Fig 4A. In Supplementary Fig S2, ROC curves are also shown for the differentiation of LUAD versus LUSC using the LUAD test score (AUC = 0.93) and LUSC test score (AUC = 0.88), respectively. At a conservative cutoff of 3.5% of the classifier’s attribution to one of the cancer subtypes (referred to as ‘indicative subtype fraction’ in Fig 4B-C), none of the control plasma were falsely predicted as lung cancer. Confusion matrices across both cohorts are shown (Figure 4B-C). Our data show that ∼72% (29/42 for UZ Gent cohort and 25/33 for UMCG cohort) of patients have sufficient circulating tumor DNA to allow for reliable non-invasive histological subtyping by our methylation profiling (Methods). In the UMCG cohort the VAF on different driver mutations was determined using the AVENIO ctDNA Expanded Kit, as a gold standard method to establish the tumor-derived fraction in the cfDNA (28). After log2 transformation, a correlation of R^2^=0.5712 between VAF and indicative subtype fraction by methylation was calculated (Fig 4D). We observed that methylation-based classification was accurate across the range of VAF/ctDNA %.

## DISCUSSION

cf-RRBS is a RRBS method that is applicable on cfDNA by effectively overcoming the challenges associated with low abundant and highly fragmented input material. MspI digestion and subsequent size-based (20-200 bp) fragment selection results in capturing the majority of promoters and CpG islands (13), uncovering most of the methylation-determined information on cellular (patho)physiology (29). With cf-RRBS, the same representative section of the epigenome is obtained, achieving a similar cost-benefit balance (i.e., ∼$100 per sample (30)) as provided by classic RRBS, rendering our method feasible for large-scale studies and routine diagnostics, especially as we have designed and implemented it for liquid handler automation. To achieve such feasible cost per test in routine diagnostics, pooling some cf-RRBS prepared samples with other NGS-based prepared samples (e.g. NIPT prepared samples) is a tested and approved option.

Until now, three other ‘whole-genome’ method concepts have been published which similarly target CpG-rich regions in highly fragmented cfDNA: oligonucleotide capture-based products (SeqCap Epi (Roche NimbleGen) and NGS methylation detection system (Twist Bioscience)) and antibody- or methylation binding domain-based enrichment (cfMeDIPSeq & T7-MBD-seq). These methods, in stark contrast to cf-RRBS’ mostly single-tube workflow, are time consuming, and require a great number of complex sample handling steps, including solid-phase reversible immobilization magnetic bead purifications, vacuum concentrator drying stages, and numerous sample transfer steps and temperature-sensitive steps. Some of these steps (e.g., drying steps) are difficult to validate in routine diagnostics workflows and the combination of all of these make automation far from trivial. We anticipate that cf-RRBS, like NIPT testing, will be much easier to implement in most molecular pathology diagnostic laboratories and cf-RRBS, as we have already observed in the laboratories with which we have collaborated for pilot clinical studies (2,17,20). cf-RRBS only relies on a few standard molecular biology enzymes and bisulfite conversion, which offers supplier versatility instead of making important diagnostic workflows dependent on a single commercial supplier of the proprietary reagents that are needed in these other methods.

It also deserves note that the methodology can directly be applied to any species’ DNA and hence applications can be anticipated in veterinary medicine, plant epigenetics etc., where cost pressures are even higher than in human diagnostics. As cf-RRBS was designed to operate on highly fragmented DNA, we have also found that the method works well to analyze the fragmented DNA extracted from formalin-fixed paraffin-embedded tissue, enabling the production of cf-RRBS methylation reference atlases directly from the rich collections of pathological archival materials (17).

The method’s application is illustrated on a relevant problem related to lung cancer diagnosis. Here, for patients with inaccessible lesions or substantial comorbidity, biopsy tissue examination might be delayed or simply not possible (31). Moreover, SCLC transformation, seen as a mechanism of therapy resistance in EGFR-driven NSCLC (25), is difficult to intercept by classic disease monitoring standards (i.e., no serial tumor biopsies possible; mostly imaging-based). Our cf-RRBS and data-derived classifier provided sufficient signal for accurate classification in 72% of the cases. For the other cases, the signal was too low, which we suspect relates to the fact that we relied on publicly available methylation reference atlases generated with a different method (Illumina arrays) resulting in only a few % of the cf-RRBS data being used (i.e. for those regions in the genome that are probed by both methods (32). Hence, as cf-RRBS also works well on FFPE archived tissue, we expect that significant further improvement is possible by establishing tumor-entity methylation reference atlases directly by cf-RRBS analysis on reference tumor tissues.

Beyond a couple of already-published oncology studies (2, 17, 20, 33), prenatal study (21) and the above-presented proof-of-principle analysis on lung cancer, clinical efforts to further explore cf-RRBS’s range of application are in process. Since the preprint of our method, other groups are also producing further derived methods that use the same concepts (34, 35). Building on ongoing rapid developments in epigenomic sequencing of tumors and non-cancerous pathologies, such as immune/inflammatory diseases (36), degenerative diseases (37), infectious diseases (38), substance abuse (39) and prenatal complications (40), cf-RRBS is poised for broad clinical utility.

## MATERIALS AND METHODS

All wet-lab and computational methods, as well as patient characteristics are included in the Online Methods.

## List of Supplementary Materials

Supplementary Methods

Figure S1 to S2

Table S1 to S5 (excel files)

## Acknowledgments

We thank Hans Van Vlierberghe and Xavier Verhelst for providing blood samples from patients with liver disease. We thank Tim Lammens and Geneviève Laureys for providing the neuroblastoma samples. We thank Jo Vandesompele, Anneleen Decock, Wim Van Criekinge and their teams for the reuse data on neuroblastoma patients. We are grateful to Wim Timens for assisting in reviewing the UMCG tumor tissue samples.

## Funding

Research Foundation Flanders 11R8616N (ADK)

Research Foundation Flanders 11B3718N (RVP)

Research Foundation Flanders 1S90621N (JDW)

Research Foundation Flanders 18B1712N (BDW)

Research Foundation Flanders 1SHGW24N (AZ)

Bijzonder Onderzoeksfonds BOF.STA.2017.0002.01 (LR)

Bijzonder Onderzoeksfonds BOF24-GOA-033-Callewaert (ADK)

Kom Op Tegen Kanker KotK_UG/2020/12444/1 (ADK)

VIB (ADK, NC)

VIB Grand Challenges project VR 2021 1712 DOC. 1492/4 (ADK, NC)

Ghent University (ADK, LR, RVP, MVdL, AZ, JDW, BDW, BM, KDR, NC)

UZ Innovation Fund (BVG)

Fees from Roche, Astrazeneca, BMS to institution UMCG (TJH)

Fees from Roche, Janssen-Cilag, Amgen, Bayer, Daiichi-Sankyo, AstraZeneca, Eli Lilly, Pfizer to institution UMCG (VdJ)

Fees from Abbott, Biocartis, AstraZeneca, Invitae, Bayer, Bio-Rad, Roche, Agena Bioscience, CC Diagnostics, MERCK, Boehringer Ingelheim, Novartis, BMS, Eli Lilly, Amgen, lllumina, Janssen Cilag, Astellas Pharma, GSK, Sinnovisionlab, Sysmex, Seracare, cieBOD to institution UMCG (ES)

## Author contributions

Conceptualization: ADK, LR, RVP, JVD, NC

Methodology: ADK, LR, RVP, JVD, NC

Software: ADK, LR, RVP, SVdV, BVG

Validation: ADK, LR, RVP, MVdL, JDW, PVdL, MZ, VdJ, TJH, ES

Formal analysis: ADK, LR, RVP, MVdL

Investigation: ADK, LR, RVP, MVdL

Resources: KV, ID, VS, UH, FD, LF, KC, PVdL, MZ, VdJ, TJH, ES

Data curation: ADK, LR, RVP, SVdV, MVdL, JDW, PVdL, MZ, VdJ, TJH, ES

Writing – original draft: ADK, LR, NC

Writing – review & editing: ADK, LR, RVP, MVdL, SVdV, PVdL, BVG, AZ, KV, ID, VS, UH, FD, LF, KC, JDW, MZ, VdJ, TJH, BDW, BM, KDR, ES, JVD, NC

Visualization: ADK, LR, RVP, SVdV, NC Supervision: BDW, BM, KDR, ES, JVD, NC

Project administration: ADK, LR, RVP, MVdL

Funding acquisition: BDW, BM, KDR, ES, JVD, NC

## Competing interests

A.D.K. and N.C. are listed as inventors in patent application PCT/EP2017/056850: MEANS AND METHODS FOR AMPLIFYING NUCLEOTIDE SEQUENCES related to the methods disclosed in this manuscript.

## Data and materials availability

The data, code and materials underlying this article are available within the main text and its supplementary materials. Code is also available upon publication on https://github.com/AndriesDeKoker/cfRRBS_paper. Fastq-files containing the methylation data are available upon publication in EGA under accession number EGAD50000001608.

## Notes

### Summary of Updates

Correction of spelling errors across the whole text Adding information about data access on github and EGA Correction in data table

## References (S1–S14)

1. Guo S, Diep D, Plongthongkum N, Fung HL, Zhang K, Zhang K. Identification of methylation haplotype blocks aids in deconvolution of heterogeneous tissue samples and tumor tissue-of-origin mapping from plasma DNA. Nat Genet. 49, 635–642 (2017).

2. Walters S, Maringe C, Coleman MP, Peake MD, Butler J, Young N, et al. Lung cancer survival and stage at diagnosis in Australia, Canada, Denmark, Norway, Sweden and the UK: a population-based study, 2004–2007. Thorax. 68, 551–564 (2013).

3. Weber S, van der Leest P, Donker HC, Schlange T, Timens W, Tamminga M, et al. Dynamic Changes of Circulating Tumor DNA Predict Clinical Outcome in Patients With Advanced Non–Small-Cell Lung Cancer Treated With Immune Checkpoint Inhibitors. JCO Precis Oncol. 5, 1540–1553 (2021).

4. Iorio F, Knijnenburg TA, Vis DJ, Bignell GR, Menden MP, Schubert M, et al. A Landscape of Pharmacogenomic Interactions in Cancer. Cell. 166, 740–754 (2016).

5. Köster J, Rahmann S. Snakemake-a scalable bioinformatics workflow engine. Bioinforma Oxf Engl. 34, 2520–2522 (2018).

6. Martin M. Cutadapt removes adapter sequences from high-throughput sequencing reads. EMBnet.journal. 17, 10–12 (2011).

7. Krueger F, Andrews SR. Bismark: a flexible aligner and methylation caller for Bisulfite-Seq applications. Bioinforma Oxf Engl. 27, 1571–1572 (2011).

8. Langmead B, Salzberg SL. Fast gapped-read alignment with Bowtie 2. Nat Methods. 9, 357–359 (2012).

9. Zerbino DR, Johnson N, Juetteman T, Sheppard D, Wilder SP, Lavidas I, et al. Ensembl regulation resources. Database J Biol Databases Curation. 2016, (2016).

10. Moss J, Magenheim J, Neiman D, Zemmour H, Loyfer N, Korach A, et al. Comprehensive human cell-type methylation atlas reveals origins of circulating cell-free DNA in health and disease. Nat Commun. 9, (2018).

11. Chatterjee A, Stockwell PA, Rodger EJ, Morison IM. Comparison of alignment software for genome-wide bisulphite sequence data. Nucleic Acids Res. 40, (2012).

12. Quinlan AR, Hall IM. BEDTools: a flexible suite of utilities for comparing genomic features. Bioinforma Oxf Engl. 26, 841–842 (2010).

13. Ritz C, Baty F, Streibig JC, Gerhard D. Dose-Response Analysis Using R. PLoS ONE 10, (2015).

14. Hand DJ, Till RJ. A Simple Generalisation of the Area Under the ROC Curve for Multiple Class Classification Problems. Mach Learn. 45, 171–186 (2001).

## References and Notes

1. Kulis M, Esteller M. DNA Methylation and Cancer. Advances in Genetics 70, 27–56 (2010).

2. Van Paemel R, De Koker A, Vandeputte C, van Zogchel L, Lammens T, Laureys G, et al. Minimally invasive classification of paediatric solid tumours using reduced representation bisulphite sequencing of cell-free DNA: a proof-of-principle study. Epigenetics 16, 196–208 (2020).

3. Kang S, Li Q, Chen Q, Zhou Y, Park S, Lee G, et al. CancerLocator: non-invasive cancer diagnosis and tissue-of-origin prediction using methylation profiles of cell-free DNA. Genome Biol. 18, (2017).

4. Li W, Li Q, Kang S, Same M, Zhou Y, Sun C, et al. CancerDetector: ultrasensitive and non-invasive cancer detection at the resolution of individual reads using cell-free DNA methylation sequencing data. Nucleic Acids Res. 46, (2018).

5. Moss J, Magenheim J, Neiman D, Zemmour H, Loyfer N, Korach A, et al. Comprehensive human cell-type methylation atlas reveals origins of circulating cell-free DNA in health and disease. Nat Commun. 9, (2018).

6. Dedeurwaerder S, Defrance M, Calonne E, Denis H, Sotiriou C, Fuks F. Evaluation of the Infinium Methylation 450K technology. Epigenomics 3, 771–784 (2011).

7. Zhou W, Laird PW, Shen H. Comprehensive characterization, annotation and innovative use of Infinium DNA methylation BeadChip probes. Nucleic Acids Res. 45, (2017).

8. Meissner A, Gnirke A, Bell GW, Ramsahoye B, Lander ES, Jaenisch R. Reduced representation bisulfite sequencing for comparative high-resolution DNA methylation analysis. Nucleic Acids Res. 33, 5868–5877 (2005).

9. Yong WS, Hsu FM, Chen PY. Profiling genome-wide DNA methylation. Epigenetics Chromatin 9, (2016).

10. Deaton AM, Bird A. CpG islands and the regulation of transcription. Genes Dev. 25 1010–1022 (2011).

11. Wendt J, Rosenbaum H, Richmond TA, Jeddeloh JA, Burgess DL. Targeted Bisulfite Sequencing Using the SeqCap Epi Enrichment System. Methods Mol Biol., 383–405 (2018).

12. Shen SY, Burgener JM, Bratman SV, De Carvalho DD. Preparation of cfMeDIP-seq libraries for methylome profiling of plasma cell-free DNA. Nat Protoc. 14, 2749–2780 (2019).

13. Gu H, Smith ZD, Bock C, Boyle P, Gnirke A, Meissner A. Preparation of reduced representation bisulfite sequencing libraries for genome-scale DNA methylation profiling. Nat Protoc. 6, 468–481 (2011).

14. Kostenko E, Chantraine F, Vandeweyer K, Schmid M, Lefevre A, Hertz D, et al. Clinical and Economic Impact of Adopting Noninvasive Prenatal Testing as a Primary Screening Method for Fetal Aneuploidies in the General Pregnancy Population. Fetal Diagn Ther. 45, 413–423 (2019).

15. Rossi R, Montecucco A, Ciarrocchi G, Biamonti G. Functional characterization of the T4 DNA ligase: a new insight into the mechanism of action. Nucleic Acids Res. 25, 2106–2113 (1997).

16. Jiang P, Lo YMD. The Long and Short of Circulating Cell-Free DNA and the Ins and Outs of Molecular Diagnostics. Trends Genet. 32, 360–371 (2016).

17. De Wilde J, Van Paemel R, De Koker A, Roelandt S, Van de Velde S, Callewaert N, et al. A Fast, Affordable, and Minimally Invasive Diagnostic Test for Cancer of Unknown Primary Using DNA Methylation Profiling. Lab Investig J Tech Methods Pathol. 104, (2024).

18. Zerbino DR, Wilder SP, Johnson N, Juettemann T, Flicek PR. The Ensembl Regulatory Build. Genome Biol. 16, (2015).

19. Taryma-Leśniak O, Sokolowska KE, Wojdacz TK. Current status of development of methylation biomarkers for in vitro diagnostic IVD applications. Clin Epigenetics. 12, (2020).

20. Cornelli L, Van Paemel R, Ferro dos Santos MR, Roelandt S, Willems L, Vandersteene J, et al. Diagnosis of pediatric central nervous system tumors using methylation profiling of cfDNA from cerebrospinal fluid. Clin Epigenetics. 16, (2024).

21. Baetens M, Van Gaever B, Deblaere S, De Koker A, Meuris L, Callewaert N, et al. Advancing diagnosis and early risk assessment of preeclampsia through noninvasive cell-free DNA methylation profiling. Clin Epigenetics. 16, (2024).

22. Chan BA, Hughes BGM. Targeted therapy for non-small cell lung cancer: current standards and the promise of the future. Transl Lung Cancer Res. 4, (2015).

23. Scagliotti GV, Parikh P, von Pawel J, Biesma B, Vansteenkiste J, Manegold C, et al. Phase III study comparing cisplatin plus gemcitabine with cisplatin plus pemetrexed in chemotherapy-naive patients with advanced-stage non-small-cell lung cancer. J Clin Oncol Off J Am Soc Clin Oncol. 26, 3543–3551 (2008).

24. Huang CY, Ju DT, Chang CF, Muralidhar Reddy P, Velmurugan BK. A review on the effects of current chemotherapy drugs and natural agents in treating non–small cell lung cancer. BioMedicine. 7, (2017).

25. Oser MG, Niederst MJ, Sequist LV, Engelman JA. Transformation from non-small-cell lung cancer to small-cell lung cancer: molecular drivers and cells of origin. Lancet Oncol. 16, 165–172 (2015).

26. Horvath S. DNA methylation age of human tissues and cell types. Genome Biol. 14, (2013).

27. Heyn H, Li N, Ferreira HJ, Moran S, Pisano DG, Gomez A, et al. Distinct DNA methylomes of newborns and centenarians. Proc Natl Acad Sci. 109, 10522–10527 (2012).

28. Weber S, van der Leest P, Donker HC, Schlange T, Timens W, Tamminga M, et al. Dynamic Changes of Circulating Tumor DNA Predict Clinical Outcome in Patients With Advanced Non–Small-Cell Lung Cancer Treated With Immune Checkpoint Inhibitors. JCO Precis Oncol. 5, 1540–1553 (2021).

29. Bock C. Epigenetic biomarker development. Epigenomics. 1, 99–110 (2009).

30. Boyle P, Clement K, Gu H, Smith ZD, Ziller M, Fostel JL, et al. Gel-free multiplexed reduced representation bisulfite sequencing for large-scale DNA methylation profiling. Genome Biol. 13, (2012).

31. Stokstad T, Sørhaug S, Amundsen T, Grønberg BH. Medical complexity and time to lung cancer treatment – a three-year retrospective chart review. BMC Health Serv Res. 17, (2017).

32. De Ridder K, Che H, Leroy K, Thienpont B. Benchmarking of methods for DNA methylome deconvolution. Nat Commun. 15, (2024).

33. Schoofs K, Ferro Dos Santos MR, De Wilde J, Roelandt S, Van de Velde S, Decruyenaere P, et al. Therapy response monitoring in blood plasma from esophageal adenocarcinoma patients using cell-free DNA methylation profiling. Sci Rep. 14, (2024).

34. Stackpole ML, Zeng W, Li S, Liu CC, Zhou Y, He S, et al. Cost-effective methylome sequencing of cell-free DNA for accurately detecting and locating cancer. Nat Commun. 13, (2022).

35. Heeke S, Gay CM, Estecio MR, Tran H, Morris BB, Zhang B, et al. Tumor- and circulating-free DNA methylation identifies clinically relevant small cell lung cancer subtypes. Cancer Cell. 42, 225–237 (2024).

36. Liu Y, Aryee MJ, Padyukov L, Fallin MD, Hesselberg E, Runarsson A, et al. Epigenome-wide association data implicate DNA methylation as an intermediary of genetic risk in rheumatoid arthritis. Nat Biotechnol. 31, 142–147 (2013).

37. Urdinguio RG, Sanchez-Mut JV, Esteller M. Epigenetic mechanisms in neurological diseases: genes, syndromes, and therapies. Lancet Neurol. 8, 1056–1072 (2009).

38. Pacis A, Tailleux L, Morin AM, Lambourne J, MacIsaac JL, Yotova V, et al. Bacterial infection remodels the DNA methylation landscape of human dendritic cells. Genome Res. 25, 1801–1811 (2015).

39. Cecil C a. M, Walton E, Smith RG, Viding E, McCrory EJ, Relton CL, et al. DNA methylation and substance-use risk: a prospective, genome-wide study spanning gestation to adolescence. Transl Psychiatry. 6, (2016).

40. Yuen RK, Peñaherrera MS, von Dadelszen P, McFadden DE, Robinson WP. DNA methylation profiling of human placentas reveals promoter hypomethylation of multiple genes in early-onset preeclampsia. Eur J Hum Genet. 18, 1006–1012 (2010).

