## Supplementary material for "Development of circulating cell-free DNA reduced representation bisulfite sequencing for clinical methylomics diagnostics"

**SUPPLEMENTARY METHODS** 1
1. Patients and Datasets 1
1.1. Method development samples 1
1.2. Lung cancer study cohorts 1
2. Library preparation and sequencing 2
2.1. Cell-free reduced representation bisulfite sequencing (cf-RRBS) 2
2.2. Automation of cf-RRBS on a TECAN EVO200 platform 4
2.3. Classical reduced representation bisulfite sequencing (RRBS) 4
2.4. Whole-genome bisulfite sequencing (WGBS) 5
2.5. SeqCap Epi 5
3. Extracting CpG site-level beta values 6
3.1. Sequencing data 6
3.2. Array data 6
4. Post-processing of CpG calls 6
4.1. Technical validation samples 6
4.2. Clinical classifier development samples 7
5. Paper figures and remaining analyses 8
**SUPPLEMENTARY TABLES** 9
**SUPPLEMENTARY FIGURES** 10
**REFERENCES** 12

### SUPPLEMENTARY METHODS

#### 1. Patients and Datasets

##### 1.1. Method development samples

**Liver pathology samples.** Written informed consent to participate was obtained from all patients (n=15). The inclusion of these cases for the purpose of cf-RRBS method development was approved by the institutional ethics committee at Ghent University Hospital (EC2016/0532). Within 4 days, plasma was obtained by centrifugation (without brake) of blood at 1900xg for 15 minutes at room temperature. Circulating cell-free DNA (cfDNA) was extracted from 5 mL plasma using the Quick-cfDNA™ Serum & Plasma Kit (ZymoResearch, Cat. No. D4076) following the manufacturer's instructions. The buffy coat, containing white blood cells, used to extract genomic DNA (gDNA), was pipetted in a 15mL tube and 10 times the volume of ACK lysis buffer (ThermoFisher scientific, Cat No. A1049201) was added. The tube was centrifuged at 300xg for 5 minutes and the supernatant was discarded. This was repeated twice. Lysis buffer from the Gentra Puregene cell kit plus (Qiagen, Cat No. 158788) was added to the cell pellet, which was kept at room temperature in a dark drawer until the cell mass completely dissolved. The kit claims stability of the DNA for 2 years in this lysis buffer. Within a couple of weeks, no cell mass could be observed, and the protocol was continued following the manufacturer's instructions: 'DNA Purification from Cultured Cells Using the Gentra Puregene Cell Kit' if processing 1-2 x $10^7$ cells.

**Neuroblastoma samples.** Two neuroblastoma patients were used in the technical method development stage of the project. The inclusion of these cases was approved by the institutional ethics committee at Ghent University Hospital (EC2008/104/mf). Between 4 to 10 mL of blood was drawn in a serum collection tube. Serum was obtained by leaving it at room temperature for 20 minutes. The tube was centrifuged for 10 minutes at 1000xg and the serum was separated from the blood clot. Serum was frozen at -80°C before circulating cfDNA was extracted using the QIAamp Circulating Nucleic Acid Kit (Qiagen, Cat. No. 55114) following the manufacturer's instructions.

Publicly available RRBS data on cfDNA collected from healthy individuals (1) were downloaded from Gene Expression Omnibus (GSE79279).

##### 1.2. Lung cancer study cohorts

**Cohort used for lung cancer cf-RRBS classifier development and testing**. Between January 2016 and June 2019, advanced stage lung cancer patients were recruited at Ghent University Hospital and AZ Delta Roeselare. Written informed consent to participate was obtained from all patients (n=42; Table S4). The inclusion of these cases was approved by the institutional ethics committee at Ghent University Hospital (EC2015/1468). Histopathological diagnostic classification of lung cancer subtypes was executed according to the 2016 World Health Organization's guidelines. Liquid biopsies were collected at initial diagnosis, progression or relapse. An advanced stage cohort was clinically most relevant since approximately 70-75% of all lung cancer cases are diagnosed as advanced stage diseases (2) (stage III and IV). All cases were reviewed by an independent expert pathologist (Jo Van Dorpe).

**Cohort used for lung cancer cf-RRBS classifier validation.** Advanced stage lung cancer patients were recruited at University Medical Center Groningen. All patients provided written consent (n=33, Table S5). The ISOcertified UMCG biobank initiative (9001:2008 Healthcare) was approved by the medical ethics committee of the University Medical Center Groningen (No. 2010/109) and made available to this study. Histopathological diagnostic classification was executed according to the 2016 World Health Organization's guidelines. All included cases were reviewed by expert pathologists at UMCG. Circulating cfDNA samples derived from plasma including information of the VAF on different driver mutations as determined using the AVENIO ctDNA Expanded NGS assay as reported earlier (3).

**Negative control samples**. For the development/testing of classifier, a cohort of true negative control samples (n=46, age range 23-61; Table S4) and patients with non-tumor pulmonary disease (n=9, age range 44-69; Table S4), including organizing pneumonia, amyloidosis, vasculitis, silicosis and pleuritis, were recruited at Ghent University Hospital. The inclusion of these cases was approved by the institutional ethics committee at Ghent University Hospital (EC2017/1207 and EC2015/1468). For the validation, an UMCG cohort of true negative control samples (n=30, age range 48-77; Table S5), healthy blood donors were recruited via and approved by Sanquin (IC NVT0612.01). Written informed consent to participate was obtained from all individuals.

**Sample collection procedure**. Blood samples were collected in Cell-Free DNA BCT tubes (10 mL) (Streck, La Vista) or PAXgene Blood ccfDNA tubes (10 mL) (PreAnalytiX, Hombrechtikon). Within 24 hours of collection, plasma isolation was executed by one or two consecutive (BCT) centrifugation steps, according to the manufacturer's protocol. For samples obtained at Ghent University Hospital, cfDNA extraction from 3.5 mL of plasma was performed using the Maxwell RSC ccfDNA Plasma Kit (Promega, Madison), following the manufacturer's instructions. For samples obtained at University Medical Center Groningen from 2ml plasma cfDNA extraction was performed using the QIAamp Circulating Nucleic Acid Kit (Qiagen, Hilden), following the manufacturer's instructions.

In the classification model development, we also made use of publicly available methylation data obtained with Infinium HumanMethylation 450k BeadChip arrays (Illumina), retrieved from the Cancer Genome Atlas (TCGA-LUAD, n=248 out of 475 (filter: primary_diagnosis == 'Adenocarcinoma, NOS'); TCGA-LUSC, n=370; Table S2) and from the work of Iorio et al. (SCLC, n=64) (GEO: GSE68379: Table S2) (4).

#### 2. Library preparation and sequencing

##### 2.1. Cell-free reduced representation bisulfite sequencing (cf-RRBS)

cfDNA was quantified using the FEMTO Pulse Automated Pulsed-Field CE Instrument (Agilent Technologies Inc., Santa Clara). The cf-RRBS protocol (https://www.protocols.io/view/cf-rrbs-protocol-pc6dize) was performed in a thermocycler with heated lid (105°C) (Bio-Rad Laboratories, Hercules).

2.5-10 ng (5 µL) of DNA was dephosphorylated in a 10 µL reaction by adding a 5 µL mixture consisting of 1 µL containing 0.01 ng (i.e., 0.1% w/w for a 10 ng input) of unmethylated lambdaDNA (Promega, Cat. No. D1521) (optional), 1 µL of recombinant Shrimp Alkaline Phosphatase (rSAP) (1U/µL) (New England Biolabs (NEB), Cat. No. M0371S), 1 µL of 10X CutSmart buffer (NEB, Cat. No. B7204S) and 2 µL of milliQ $H_2O$. The mixture was pipetted in 0.2 mL thin-walled PCR tubes (ThermoFisher Scientific, Cat. No. AB0622) and incubated for 1 hour at 37°C. The phosphatase was heat-inactivated for 30 minutes at 75°C.

The DNA was digested by adding 5 µL of a mixture containing 0.5 µL of MspI (20U/µL) (NEB, Cat. No. R0106S), 0.5 µL of 10X CutSmart and 4 µL of milliQ $H_2O$ and incubated for 30 minutes at 37°C. End-repair and dA-tailing was performed by adding a 10 µL-mixture of 0.5 µL of Klenow Fragment (3'->5' exo-) (5U/µL) (NEB, Cat. No. M0212S), 0.25 µL of dNTP mix (4 mM dATP, 400 µM dCTP and 400 µM dGTP) (Promega, Cat. No. U1330), 1 µL of 10X CutSmart and 8.25 µL of milliQ $H_2O$. This 25 µL-mixture was incubated for 20 minutes at 30°C, for 20 minutes at 37°C and heat-inactivated for 20 minutes at 75°C.

Next, adapters were ligated to the DNA fragments: a 10 µL-mixture containing 4 µL of 10 mM ATP (NEB, Cat. No. P0756S), 1 µL of 10 µM adapter, 1 µL of 10X CutSmart and 4 µL of milliQ $H_2O$ and a 5 µL-mixture containing 0.5 µL of T4 DNA ligase (2000 U/µL) (NEB, Cat. No. M0202M), 0.5 µL of 10X CutSmart and 4 µL of milliQ $H_2O$ was added. The adapter is identical to the NEBNext-adapter (seq: 5′-/Phos/GATCGGAAGAGCACA-CGTCTGAACTCCAGTC/ideoxyU/ACACTCTTTCCCTACACGACGCTCTTCCGATCT-3′) and was ordered at IDT, but without phosphorothioate bond at the 3'-end. Ligation was done overnight (14 hours) at 16°C and the ligase was heat-inactivated at 65°C for 10 minutes.

The next day, a mixture of 0.5 µL of exonuclease I (20 U/µL) (NEB, Cat. No. M0293S), 0.5 µL of exonuclease III (50 U/µL) (NEB, Cat. No. M0206S), 0.5 µL of exonuclease VII (10U/µL) (NEB, Cat. No. M0379S), 0.5 µL of 10X CutSmart and 3 µL of milliQ $H_2O$ was pipetted to the 40µL-reaction. This mixture was incubated for 2 hours at 37°C and heat-inactivated for 10 minutes at 80°C and 10 minutes at 95°C. As a last step before bisulfite conversion, 1 µL of Antarctic Thermolabile UDG (1U/µL) (NEB, Cat. No. M0372S), 0.5 µL of endonuclease VIII (10U/µL) (NEB, Cat. No. M0299S), 0.5 µL of 10X CutSmart and 3 µL of milliQ $H_2O$ was added and incubated for 1 hour at 37°C and heat-inactivated for 10 minutes at 75°C. The MspI/MspI-digested adapter-ligated DNA fragments were bisulfite converted using the EZ DNA Methylation-Lightning™ Kit (ZymoResearch, Cat. No. D5030) according to manufacturer's instructions. Bisulfite-converted DNA was eluted in 14 µL of milliQ $H_2O$.

A final amplification step was performed using the KAPA HiFi HotStart Uracil+ ReadyMix PCR Kit (KAPA Biosystems, Cat. No. KK2801). 15 µL of 2X polymerase mix was combined with 13.2 µL of DNA, 0.9 µL of 10 µM NEBNext universal primer (IDT) and 0.9 µL of 10 µM NEBNext index primer (IDT). The PCR protocol was: 5 minutes 95°C, 17-19x (20 seconds 98°C, 15 seconds 65°C, 45 seconds 72°C), 5 minutes 72°C, until further processing 12°C.

Libraries prepared using the cf-RRBS protocol were cleaned up by magnetic bead cleanup (CleanNA PCR) (GC biotech, Cat. No. CPCR-0050) and eluted in 0.1X TE buffer, and visualized with the Fragment Analyzer (Agilent Technologies Inc., Santa Clara), and quantified using the Qubit dsDNA HS Assay Kit (ThermoFisher scientific, Cat. No. Q32851). Based on the concentration value obtained, the libraries were pooled and were sequenced on a NextSeq500 instrument (Illumina, San Diego), performing a PE75 run using 5% phiX or a NovaSeq S1 or S2 instrument (Illumina, San Diego), performing a PE50 or PE100 run using 3% or 1% phiX.

#### 2.2. Automation of cf-RRBS on a TECAN EVO200 platform

Instead of using thin-walled PCR tubes, samples are now put in 250 µL, semi-skirted twin.tec® PCR plate 96 LoBind® (Eppendorf, Cat. No. 0030129504).

To allow the addition of standardized volumes, DNA should be dissolved at equal volume. In practice, equal amounts of DNA are pipetted into the wells and all are filled to the volume of the sample with the largest volume which was needed to get an equal amount of DNA. When every well is at equal volume, the whole 96-well plate is placed in a SpeedVac concentrator to lower the volume to 5 µL.

Mastermixes are prepared to be added as 5 µL spike-ins to facilitate accurate and robust robotic pipetting, which required that the end-repair and A-tailing mix was changed to be 0.5 µL of Klenow Fragment (3'-->5' exo-) (5U/µL), 0.2 µL of dNTP mix (4 mM dATP, 400 µM dCTP and 400 µM dGTP), 0.5 µL of 10X CutSmart and 3.8 µL of milliQ $H_2O$. Also, the combination of the adapter mixture and T4-ligase mixture was adapted to a 5 µL-mixture containing 1 µL of 10 µM adapter, 0.5 µL of 10X CutSmart and 3.5 µL of milliQ $H_2O$ and a 5 µL-mixture containing 0.5 µL of T4 DNA ligase (2000 U/µL), 0.5 µL of 10X CutSmart, 3 µL of 10 mM ATP, and 1 µL of milliQ $H_2O$. Recently exonuclease VIII, truncated (NEB, Cat. No. M0545S) has been made commercially available and we used it to replace exonuclease VII. Exonuclease VIII, truncated is a faster working enzyme. Incubation can now be done for 1 hour at 37°C and heat-inactivated for 30 minutes at 80°C.

All incubations were performed in an off-deck thermocycler. As we did not have proper plate seals available, the whole plate was recapped by 8-well strips. Mind that reusing these strips throughout the whole protocol is possible, but one has to avoid mixing up strips for different plate columns.

For the bisulfite conversion, the automation-friendly EZ-96 DNA Methylation-Lightning MagPrep was used according to manufacturer's instructions. However, all plastics which are included in the kit were necessarily exchanged for LoBind versions: Deepwell Plate 96/1000µL, DNA LoBind® (Eppendorf, Cat. No. 0030503201).

During library amplification the NEBNext dual indexing primer system, allowing 96-plex multiplexing, was used.

#### 2.3. Classical reduced representation bisulfite sequencing (RRBS)

cfDNA was quantified using the FEMTO Pulse Automated Pulsed-Field CE Instrument (Agilent Technologies Inc., Santa Clara) and gDNA using the Nanodrop (ThermoFisher Scientific, Waltham).

10 ng of DNA and 0.1 % w/w of unmethylated lambdaDNA was digested for 30 minutes at 37°C with 10 units of MspI in a 20 µL CutSmart-buffered reaction. Next, 0.9 µL of 1 mM dNTP mix (Promega, Cat. No. U1511), 2.5 units of Klenow Fragment (3' to 5' exo-), 0.5 µL 10x CutSmart buffer and 3.1 µL of milliQ $H_2O$ was added and incubated for 20 minutes at 30°C and 20 minutes at 37°C. The reaction was stopped by heat-inactivation during 30 minutes at 75°C.

Before purification by Nucleospin gel and PCR cleanup (Macherey-Nagel, Cat. No. 740609.250), 400 ng of carrier DNA was added, using 4 µg of unmethylated lambda DNA which was previously incubated for 1 hour at 37°C with 20 units of MspI and 1 unit of rSAP in a CutSmart-buffered 20 µL reaction. The reaction was stopped by heat-inactivation for 30 minutes at 75°C.

A completely methylated adapter was prepared. The adapter was ordered at IDT as two oligos (5'-ACACTCTTTCCCTACACGACGCTCTTCCGATCT-3' and 5'-/Phos/GATCGGAAGAGCACACGTCTGA-ACTCCAGTCAC-3'). To allow for adapter hybridization, 10 µL of both 100 µM oligo solutions were mixed with 2.5 µL 10X T4 DNA ligase buffer, 2.5 µL milliQ $H_2O$ and heated for 1 minute at 95°C, then cooled down to room temperature at 0.1°C/s in a thermocycler.

After cleanup of the Klenow (3' to 5' exo-) reaction, the eluate (14 µL) was mixed with 1 µL of 40 µM methylated adapter, 2 µL of 10X T4 DNA ligase buffer (NEB, Cat. No. B0202S) and 2 µL of milliQ $H_2O$. Then, 400 units of T4 DNA ligase was added. The reaction was incubated overnight at 16°C and stopped by heat-inactivation for 20 minutes at 65°C. The ligation mixture was loaded on a 3% agarose gel and ran for 1.5 hours at 105 V. The gel was imaged after Ethidium Bromide staining and the gel section corresponding to 85-285 bp was cut out. DNA was extracted from the gel using the Nucleospin gel and PCR cleanup kit and eluted in 20 µL of milliQ $H_2O$.

The DNA was bisulfite converted using the EZ DNA Lightning kit following the manufacturer's instructions. The bisulfite-treated sample (14µL) was then mixed with 15 µL of KAPA HiFi HotStart Uracil+ ReadyMix PCR Kit and 0.9 µL of 10 µM of NEBNext universal primer and 0.9 µL of 10µM of NEBNext index primer, both ordered at IDT, and amplified according to the PCR protocol: 5 minutes 95°C, 20x (20 seconds 98°C, 15 seconds 65°C, 45 seconds 72°C), 5 minutes 72°C, terminal cool to 4°C.

Libraries were cleaned by magnetic bead cleanup (CleanNA PCR) and eluted in 0.1X TE buffer, and visualized with the Fragment Analyzer (Agilent Technologies Inc., Santa Clara), and quantified using the Qubit dsDNA HS Assay Kit. Based on the concentration value obtained, the libraries were pooled and were sequenced on a NextSeq500 instrument (Illumina, San Diego), performing a PE75 run using 5% phiX or a NovaSeq S1 or S2 instrument (Illumina, San Diego), performing a PE50 or PE100 run using 3% or 1% phiX.

#### 2.4. Whole-genome bisulfite sequencing (WGBS)

The WGBS protocol was performed by the genome analysis service facility, NXTGNT (Ghent, Belgium), using 50 ng of DNA extracted from blood serum of neuroblastoma patients. The Accel-NGS methyl-seq DNA library kit for Illumina with associated indexing kit A (Swift Biosciences, Cat. No. 30024 and 36024) was used according to manufacturer's instructions. The EZ DNA Methylation-Lightning™ Kit was used for bisulfite conversion. Sequencing was performed on a HiSeq2500 (Illumina, San Diego) at BGI Genomics (Shenzhen, China) using one lane per sample, performing a PE125 run, V4 reagent.

#### 2.5. SeqCap Epi

The Accel-NGS Methyl-Seq DNA library kit with associated indexing kit A was combined with the SeqCap Epi enrichment system (Roche Nimblegen, Cat. No. 07138881001). As an internal standard, unmethylated lambda DNA was sheared to ±150bp with a S202 Focused-Ultrasonicator (Covaris, Woburn) and spiked-in before library preparation (e.g., 0.1% w/w for a 10 ng input). Library preparation was performed on 2.5-10 ng of input DNA following manufacturer's instructions, taking into account their proposed adaptations when working with DNA and combining with the NimbleGen SeqCap Epi Hybridization capture probes. To cut costs, before hybridization capture, six libraries were pooled (166 ng/sample) and also 1 nmol of all appropriate SeqCap HE Index Blocking Oligo's were added. Sequencing was performed on a HiSeq4000 (Illumina, San Diego) using two lanes, performing a PE150 run, V4 reagent with 10% phiX spike-in or a NovaSeq S2 (Illumina, San Diego), performing a PE100 run with 1% phiX spike-in.

#### 3. Extracting CpG site-level beta values

##### 3.1. Sequencing data

A FreeBSD-configured storage server was used to demultiplex raw reads from cf-RRBS data by bcl2fastq (v2.20.0.422; Illumina, San Diego). Public sequencing data, in form of fastq files, were downloaded from the given sources. Using the high-performance computing infrastructure of Ghent University, the raw reads were processed by a custom snakemake (v5.2.4) (5) pipeline:

- https://github.com/rmvpaeme/cfRRBS_tube_study/tree/master/code/preprocessing
- Trim Galore (v0.6.0) (https://www.bioinformatics.babraham.ac.uk/projects/trim_galore/) was used to trim reads (fastq files), internally using Cutadapt (v1.18) (6), using a Phred score limit of minimum 20; a maximum trimming error rate of 0.1; a minimum required detected adapter of 1 bp; and a minimum required read length of 20 bp. Variable parameters were:
    - **(cf-)RRBS**. *--rrbs --three_prime_clip_R1 1 --three_prime_clip_R2 1 --clip_R1 3 --clip_R2 3*
    - **SCE**. *--clip_R1 10 --clip_R2 10 --three_prime_clip_R1 10 --three_prime_clip_R2 10*
    - **WGBS**. *--clip_R1 18 --clip_R2 18 --three_prime_clip_R1 2 --three_prime_clip_R2 2*
- Mapping to the human reference genome (GRCh38, without alt chromosomes) was executed by Bismark (7) (v0.20.1), which internally uses Bowtie2 (v2.3.4) (8) to generate bam files. For WGBS data, a deduplication step (given it concerns non-targeted data) was performed using Bismark's *deduplicate_bismark* functionality.
- Optionally: filter_non_conversion -p --percentage_cutoff 90 --minimum_count 3
- Methylation calling was finally performed by Bismark's *bismark_methylation_extractor* package.

##### 3.2. Array data

For LUAD and LUSC, datasets were downloaded using the TCGAbiolinks R library (data release v43.0).

For SCLC, raw signal intensities were obtained. Methylation calls were deduced using the R package openSesame (v1.20.0) by its function *getBetas* under default parameters, performing background correction, dye-bias normalization and calculation of beta values.

#### 4. Post-processing of CpG calls

##### 4.1. Technical validation samples

To increase robustness and decrease computational complexness, exclusively CpG sites covered by sequencing five or more times were kept. Sites were mapped to CpG islands defined according to the human reference sequence (*cpgIslandExt* from UCSC, updated 08-2018) and regulatory elements (regulatory build from Ensembl, release 103, updated 02-2021) (9). For the regulatory elements, unannotated ChIP-Seq transcription factor binding sites and DNase I hypersensitivity open chromatin regions were grouped together as one category. Since CTCF binding sites often (partly) overlap with other elements, exclusively unique CTCF binding site stretches were inferred to form a new 'CTCF binding site-only' category. The beta value of grouped CpG sites (i.e., islands and regulatory elements) was calculated as the mean of individual CpG sites.

#### 4.2. Clinical classifier development samples

**CpG clustering**. Instead of using predefined clusters, such as CpG islands or regulatory elements, new clusters were defined optimized for cf-RRBS in order to reduce data complexity (~3-4M data points to ~500k data points per sample) and obtain robust and biologically relevant modeling features without losing data. The following *adhoc* algorithm was used, making use of all cf-RRBS samples of the 'clinical classifier development samples':

- CpG methylation calls not observed in 50% or more of the healthy control data set are removed; clustering is executed on the remaining CpG methylation calls (*~CpG set X*).
- The maximum allowed length between two CpGs in this *set X* is 50 nucleotides in order for them to be considered to be in one cluster.
- Each cluster carries at least two CpGs
- The beta value of a cluster equals the mean of the beta values of its enclosed CpGs.

**Mapping of public data to clusters**. The public lung cancer tissue methylation data was mapped to these clusters by adhering following rules:

- **Arrays**. CpG sites were directly mapped to the clusters. Cluster beta values were calculated as the mean of their enclosed CpG sites. When no CpG calls were present, the cluster was defined as 'missing' in the corresponding public sample.

**Predictive modeling – feature selection**. A two-step procedure was used to select features (CpG clusters) with high discriminative power:

- Within the set of in-house healthy negative controls sequenced by cf-RRBS (n=46; Table S4), unstable clusters were removed (standard deviation beta value >1%). Doing this, we aimed at obtaining a subset of clusters that is unrelated to age or environmental covariables and thus constant in a healthy control population.
- For each subtype of interest (LUAD, LUSC and SCLC), the 250 most divergent clusters from the above stable cluster healthy negative control data set, according to the mean absolute difference with public methylation data (LUAD, n=248 of 475 (filter: primary_diagnosis == 'Adenocarcinoma, NOS'); LUSC, n=370; and SCLC, n=64; Table S2), were selected as final features.

**Predictive modeling – deconvolution**. Lung cancer subtype modeling was performed by meth_atlas (10), which executes deconvolution (i.e., non-negative least squares) to estimate the relative contribution of each reference entity (healthy background, cancer subtypes, …) in a given liquid biopsy. In the context of this study, four possible contributions/classes were allowed and used for classifier training: negative control liquid methylation patterns; and LUAD, LUSC and SCLC tissue methylation patterns. The training atlas contains average beta values per class of the most diagnostic discriminating methylation cluster features selected above. Finally, we tested the classifier model on cf-RRBS liquid biopsies for the in-house cohort (LUAD, n=18; LUSC, n=12; SCLC, n=12; non-tumor lung disease negative control, n=9 and healthy control, n=46; Table S4). Subsequently, this data-analysis and

diagnostic classification method was independently validated in a separate cohort from a different hospital (LUAD, n=22; LUSC, n=11 and healthy control, n=30; Table S5).
From this first analysis, we determined that at least 3,5% of the cf-RRBS data needed to be assigned in this way by the algorithm to a particular deconvolution class, for this to be sufficient for diagnosis.

#### 5. Paper figures and remaining analyses

All plots were made by custom scripting in R (v4.3.1).

**Capillary gel electropherogram**. The capillary gel electropherogram image was processed by ProSize (v3.0; Agilent Technologies Inc., Santa Clara) (Figure 1B).

**Target fold enrichment**. A genome-wide target file was created *in silico* using mkrrgenome (v1.23) (11), selecting GRCh38 MspI/MspI fragments sized between 20 and 200 nucleotides, translated to bed format by custom scripting. Target fold enrichments were calculated from bam files by Picard Tools (v2.18.27) (https://broadinstitute.github.io/picard/), through *CollectHsMetrics* (Figure 2A, 2C).

**Mapped reads in function of genomic target size**. The number of mapped reads in function of genomic target size was obtained by the *coverage* functionality of BEDTools (v2.29.2) (12) (Figure 2B).

**Per base fold enrichment**. Mapped read positions were extracted from bam files by GNU Awk (v4.0.2). The per base fold enrichment was calculated as the number of reads that cover a position over the average number of reads that cover any position (~the overall genome coverage) (Figure 2D).

**Pearson correlation and mean absolute error between two methylation profiles**. The positional intersection of CpG sets was considered to calculate beta value Pearson correlations and mean absolute errors (~the mathematical mean absolute difference) (Figure 3A, 3B, 3C & Figure S1).

**CpG coverage overlap**. The positional overlap in CpG calls between two methylation profiles was calculated as the number of recurrent CpG calls (intersection) over the total number of CpG calls in the sample with the lowest sequencing depth—this to minimize coverage difference bias (Figure 3C).

**Log regression fitting**. Log regression fitting was executed by the R package drc (v3.0-1) (13) (Figure 3E).

**Multi-class receiver operating characteristic analysis (14)**. Analyses to determine the areas under the operating characteristic curves (AUROCs) were executed to obtain cutoff-free classification performance statistics using the pROC and ROCit R-library (v1.18.5 & v2.1.1). For each case of differential diagnosis (i.e., healthy vs non-healthy, SCLC vs NSCLC, LUAD vs LUSC), the maximal predicted fractions of the corresponding subtype (i.e., healthy fraction for the healthy vs non-healthy problem, SCLC fraction for the SCLC vs NSCLC problem, LUAD or LUSC fraction for the LUAD vs LUSC problem) was used directly to execute a receiver operating characteristic analysis (Figure 4A & Figure S2).

#### SUPPLEMENTARY TABLES

Table S1. **Technical method development samples**. In-house sequenced data is grouped separately from publicly available data. The number of patients used in each setting is provided in the first column. Each cell indicates which type(s) of DNA source (cfDNA or gDNA) was/were analyzed by which technique (i.e., column name). Analyses shown in green were excluded from the general cf-RRBS performance-related investigations.

Table S2. **IDs of the public LUAD, LUSC and SCLC data used**

Table S3A. **Positional CpG clusters optimized for cf-RRBS, sorted according to position**. The gene associated with the nearest transcription start site, and nearest regulatory elements, are provided per cluster. When a cluster covers more than one transcription start site or regulatory element, these are given using a comma-separated list.

Table S3B. **Selected CpG clusters for use in lung cancer classifier, sorted according to genomic position**. Average beta values are depicted for the four considered classes. The nearest transcription start site (where the distance is emphasized using a grey-to-white color spectrum, mapped to values of 0 until 1000 nucleotides), corresponding genes, and nearest regulatory elements, are provided per cluster. When a cluster covers more than one transcription start site or regulatory element, these are given using a comma-separated list.

Table S4. **In-house lung cancer patient and healthy control population samples with key characteristics**. Abbreviations: CPR, central pathology review; WBRT, whole brain radiation therapy; *, only cytological specimen present during pathological examination.

Table S5. **Validation lung cancer patient and healthy control population samples with key characteristics**. Abbreviations: CPR, central pathology review.

#### SUPPLEMENTARY FIGURES

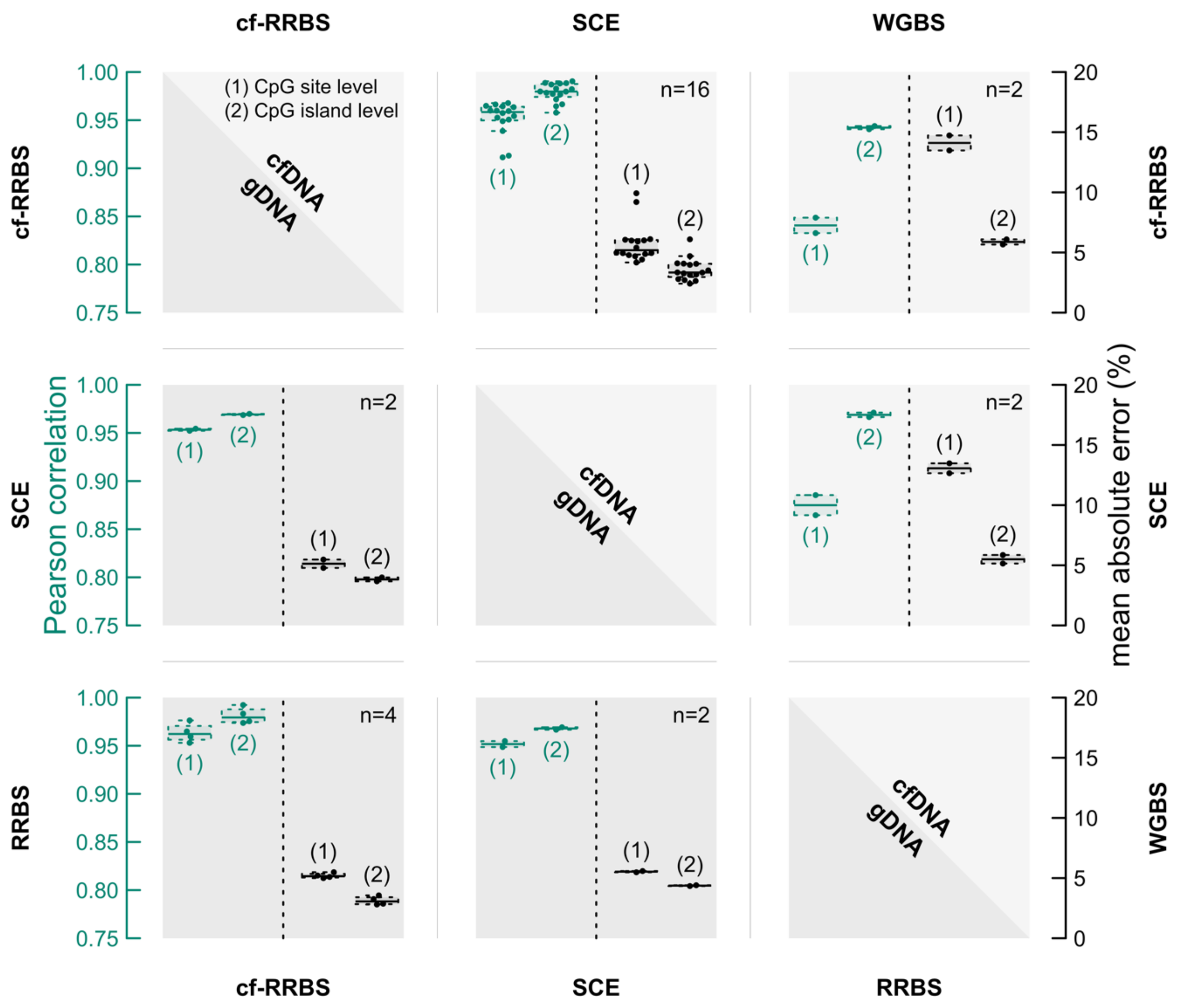


Figure S1. **Matrix visualizing pairwise concordance between techniques**. DNA types include cfDNA (upper right-hand half) and gDNA (lower left half). Each cell contains four swarm/box plot combinations, demonstrating concordance between the methods shown as matrix row/column names. The first two plots (CpG site, CpG island; green) indicate the Pearson correlation, with corresponding axes at the left (green); the remaining two plots (CpG site, CpG island; black) depict the mean absolute error, with corresponding axes at the right-hand side (black).

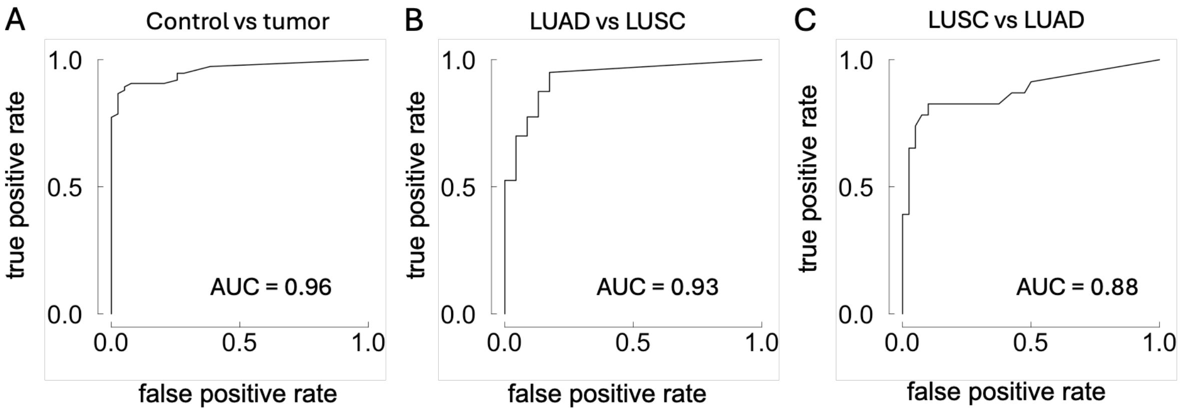


Figure S2. **Receiver operating characteristic analyses for lung cancer differential diagnosis**. AUCs are given as performance measures.

### REFERENCES


1. Guo S, Diep D, Plongthongkum N, Fung HL, Zhang K, Zhang K. Identification of methylation haplotype blocks aids in deconvolution of heterogeneous tissue samples and tumor tissue-of-origin mapping from plasma DNA. Nat Genet. 49, 635–642 (2017).

2. Walters S, Maringe C, Coleman MP, Peake MD, Butler J, Young N, et al. Lung cancer survival and stage at diagnosis in Australia, Canada, Denmark, Norway, Sweden and the UK: a population-based study, 2004–2007. Thorax. 68, 551–564 (2013).

3. Weber S, van der Leest P, Donker HC, Schlange T, Timens W, Tamminga M, et al. Dynamic Changes of Circulating Tumor DNA Predict Clinical Outcome in Patients With Advanced Non–Small-Cell Lung Cancer Treated With Immune Checkpoint Inhibitors. JCO Precis Oncol. 5, 1540–1553 (2021).

4. Iorio F, Knijnenburg TA, Vis DJ, Bignell GR, Menden MP, Schubert M, et al. A Landscape of Pharmacogenomic Interactions in Cancer. Cell. 166, 740–754 (2016).

5. Köster J, Rahmann S. Snakemake-a scalable bioinformatics workflow engine. Bioinforma Oxf Engl. 34, 2520–2522 (2018).

6. Martin M. Cutadapt removes adapter sequences from high-throughput sequencing reads. EMBnet.journal. 17, 10–12 (2011).

7. Krueger F, Andrews SR. Bismark: a flexible aligner and methylation caller for Bisulfite-Seq applications. Bioinforma Oxf Engl. 27, 1571–1572 (2011).

8. Langmead B, Salzberg SL. Fast gapped-read alignment with Bowtie 2. Nat Methods. 9, 357–359 (2012).

9. Zerbino DR, Johnson N, Juetteman T, Sheppard D, Wilder SP, Lavidas I, et al. Ensembl regulation resources. Database J Biol Databases Curation. 2016, (2016).

10. Moss J, Magenheim J, Neiman D, Zemmour H, Loyfer N, Korach A, et al. Comprehensive human cell-type methylation atlas reveals origins of circulating cell-free DNA in health and disease. Nat Commun. 9, (2018).

11. Chatterjee A, Stockwell PA, Rodger EJ, Morison IM. Comparison of alignment software for genome-wide bisulphite sequence data. Nucleic Acids Res. 40, (2012).

12. Quinlan AR, Hall IM. BEDTools: a flexible suite of utilities for comparing genomic features. Bioinforma Oxf Engl. 26, 841–842 (2010).

13. Ritz C, Baty F, Streibig JC, Gerhard D. Dose-Response Analysis Using R. PLoS ONE 10, (2015).

14. Hand DJ, Till RJ. A Simple Generalisation of the Area Under the ROC Curve for Multiple Class Classification Problems. Mach Learn. 45, 171–186 (2001).
